# Discrimination of finger movements by magnetomyography with optically pumped magnetometers

**DOI:** 10.1101/2023.03.27.534368

**Authors:** Antonino Greco, Sangyeob Baek, Thomas Middelmann, Carsten Mehring, Christoph Braun, Justus Marquetand, Markus Siegel

**Author notes:** These authors contributed equally (Antonino Greco, Sangyeob Baek). shared senior authorship (Justus Marquetand, Markus Siegel).

## Abstract

Optically pumped magnetometers (OPM) are quantum sensors that offer new possibilities to measure biomagnetic signals. In magnetomyography (MMG), compared to the current standard surface electromyography (EMG), OPM sensors offer the advantage of contactless measurements of muscle activity. However, little is known about the relative performance of OPM-MMG and EMG, e.g. in their ability to detect and classify finger movements. To address this, we recorded simultaneous OPM-MMG and EMG of finger flexor muscles for the discrimination of individual finger movements. Using a deep learning model for movement classification, we found that both sensor modalities were able to discriminate finger movements with above 89% accuracy. Furthermore, model predictions for the two sensor modalities showed high agreement in movement detection (85% agreement; Cohen’s kappa: 0.45). Our findings show that OPM sensors can be employed for reliable, contactless discrimination of finger movements and incentivize future applications of OPM in magnetomyography.

## Introduction

Spin exchange relaxation free (SERF) optically pumped magnetometers (OPM) are quantum sensors for measuring magnetic flux signals with a sensitivity in the order of few 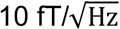. SERF-OPM are based on a zero-field resonance caused by the Zeeman-effect^1^ detected be laser spectroscopy of spin-polarized alkali metal vapor^2^. In recent years, several studies have shown the potential of OPM for measuring biomagnetic signals of the brain, heart, nerves, or muscles, opening up new opportunities for the research and application of human biomagnetism^3–9^. In particular, in addition to more traditional applications in magnetoencephalography (MEG)^5,10,11^, OPM are also increasingly utilized for studying skeletal muscles^12–14^. The spatial flexibility, small physical size (few cubic centimeters) and possibility of bi- or triaxial signal acquisition of OPM, enable the contactless investigation of muscle physiology in space and time^14^. Magnetomyography (MMG) has several general advantages in comparison to the current gold standard for non-invasive muscle studies, surface electromyography (EMG)^15,16^. In contrast to electric currents, magnetic fields are far less affected by the different tissue layers between the electromagnetic source and the skin surface, resulting in less distorted signals^12,15,17,18^. Furthermore, in contrast to MMG, EMG electrodes require contact with the skin where the presence of a charge at the electrode-skin interface creates noise voltages that can interfere with the signal^18,19^.

Magnetic flux and electric potentials originate from the same ionic currents and have comparable temporal and spectral profiles^16,20^, but the magnetic flux direction is orthogonal to the source (electric current). Newly available OPM offer up to triaxial simultaneous signal acquisition (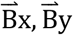, and 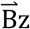) with one sensor, i.e., three-dimensional spatial information on the magnetic flux vector per sensor. The gain in spatial information per sensor in comparison to EMG could be particularly relevant for human-machine interfaces based on muscle signals, such as e.g. prosthesis control^21–23^. Considering this hypothesis and the current progress in EMG-based human-machine interfaces^24^, in the present study, we sought to investigate the potential of biomagnetic measurements as a new modality for human-machine interfaces. Specifically, we investigated and compared the ability to differentiate individual finger movements based on EMG and OPM-MMG of the finger flexor muscles.

## Methods

### Participant

A single human participant volunteered for the study. The participant of the study was male, 28 years old, 1.76 m tall, and 72 kg weight (BMI = 23.2 kg/m^2^). The participant was one of the authors and gave informed consent before participating in the study. The study was conducted in accordance with the Declaration of Helsinki and approved by the ethics committee of the University of Tübingen.

### Experimental design

The experiment was designed to record the magnetic activity of the right flexor muscles of the fingers Digit II and V (index and little finger; DII and DV). Muscle activation was measured on the forearm using simultaneous OPM-MMG and EMG (Fig. 1A-B, 4 bipolar EMG and 4 biaxial OPM channels). Simultaneously, finger movements were measured using a light sensor. The participant was instructed to flex a finger and then to return to the rest position (open hand), in three different sessions. In the first session, the participant flexed DII for 30 trials, in the second session, the participant flexed DV for 30 trials, and in the third session, the participant alternated between DII and DV for 15 trials, respectively. Every 5 seconds, an auditory cue indicated when to execute the finger movement. To measure only the movement of the designated finger, we fixated the other fingers with casts (Fig. 1A).

**Figure 1.**
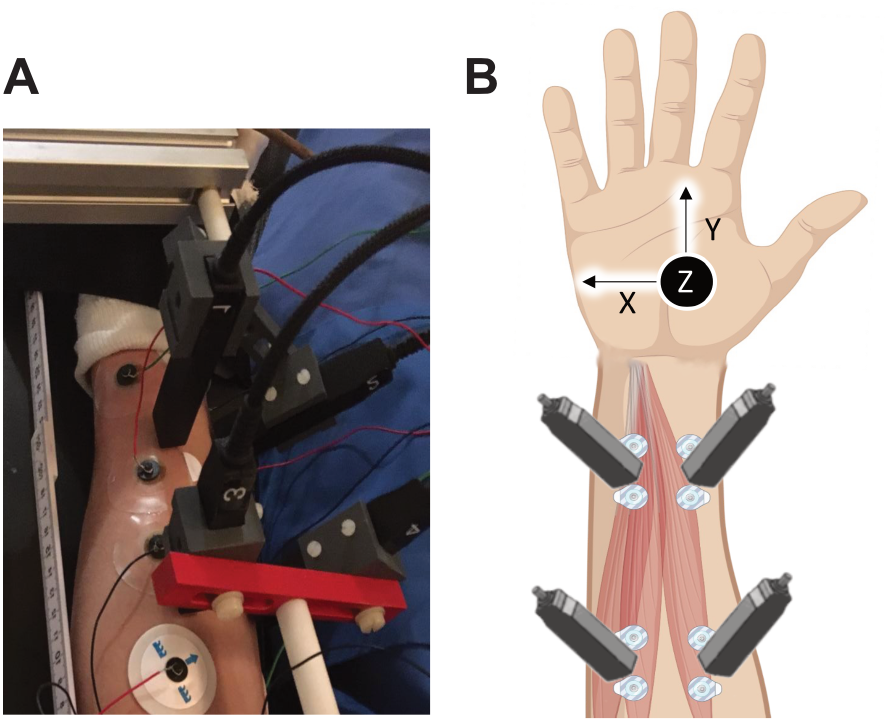
**A.** Photograph depicting the experimental settings. **B.** Conceptual representation of the sensors modalities and the location they were placed.

### Sensors

The muscle strands of the individual fingers (DII, DIV, and DV) were imaged using high-resolution muscle ultrasound (Mindray TE7, 14Mhz-linear probe) to determine the longitudinal axis of the muscles. All recordings were collected inside a magnetically shielded room (Ak3b, VAC Vacuumschmelze, Hanau, Germany). Here, 8 paramagnetic EMG surface electrodes (Conmed, Cleartrace^2^ MR-ECG-electrodes) were placed in a bipolar montage along the longitudinal axis of the muscle. A ground electrode was placed on the right shoulder. 4 biaxial OPM (QZFM-gen-1.5, QuSpin Inc., Louisville, CO, USA) were placed in between the EMG electrode pairs about 15 mm above the skin surface. The movement of the fingers was measured using a fiber optic that measured the distance between the finger and the fiber optic (Keyence Digital Fiber Sensor FS-N10).

### Data acquisition

The analog output of the OPM amplifier was recorded using the data acquisition electronics of a MEG system (CTF Omega 275, Coquitlam, BC, Canada) and the EEG channels of the MEG system were used to record the EMG. Both the OPM and EMG recordings were acquired with a sampling rate of 2343.8Hz. The employed OPMs were capable of measuring two components of the magnetic field vector: the y- and z-axis. They provided a magnetic field sensitivity of 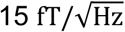 in a bandwidth of 3-135 Hz, an operating range below 200 nT, and a dynamic range of a few nanoteslas. To adapt to a non-zero magnetic background field, the sensors are equipped with internal compensation coils that can cancel magnetic background fields of up to 200 nT in the sensing hot rubidium vapor cell (cell size 3 × 3 × 3 mm).

### Data Preprocessing

Data analysis was performed using Python (Python Software Foundation, version 3.7, http://www.python.org). Data from 8 OPM channels and 4 bipolar EMG channels were demeaned and filtered using a 25-100 Hz band-pass zero-phase fourth-order butterworth infinite impulse response (IIR) filter. Line noise was filtered using a 49-51 Hz band-stop zero-phase fourth-order butterworth IIR filter. Then, we extracted the envelope of the signal by taking the absolute value of the Hilbert transformed signal and resampled the data at 200 Hz.

### Data sampling

We designed our data analysis pipeline to approximate online detection of finger movements. To this end, we aggregated all data from the three sessions and randomly sampled 100 ms windows of the signals across all channels, sensor modalities (OPM and EMG) and the finger movement signal. Crucially, the sampling procedure was nested into a cross-validation scheme (see Classification analysis section), in order to separate the training and test set at the trial level. This simulates a realtime system that scans the signal, every 100 ms, and detects finger movements. We sampled 500 windows across all the recordings for each event from the aggregate data to build the training dataset and 100 windows for the test set, where an event consisted of no motion for both fingers (DII and DV, from 1500 ms to 0 ms relative to the onset of each finger movement), DII motion and DV motion (both from 0 ms to 300 ms after the onset of each finger movement).

### Classification analysis

We performed classification analysis using a supervised learning approach using a deep convolutional neural network implemented in Tensorflow^32^. The network architecture consisted of an input layer that received batches of data consisting of matrices *T* × *C*, where *T* is the number of time points and *C* is the number of channels. Thus, the resulted input was a 3-dimensional array *N* × *T* × *C*, where *N* is the batch size that we set to 250. This tensor was passed to a residual block^25^, consisting of a point-wise (kernel size of 1 × 1) 1-dimensional convolutional layer with 128 filters, stride 1 and without the bias term and padding, followed by a batch normalization (BN)^33^ layer and a gaussian error linear unit (GELU)^34^ activation function. Then, the residual block continued with a second zero-padded 1-dimensional convolutional layer with a kernel size of 3 × 3 and *C* filters, stride 1 and without the bias term, followed by a BN layer and GELU function and concluded by adding the input array to the resulted array so far and finally applying a GELU function. After the residual block, the network architecture comprised a zero-padded 1-dimensional convolutional layer with a kernel size of 3 × 3 and 16 filters, stride 2 and with the bias term and GELU function. Then, the output of this layer was flattened and passed to a Dropout layer with a dropout rate of 0.1. From here, the output array was passed to two fully-connected (FC) layers with *Z* units and GELU as activation function (here, *Z*=100), both followed by a Dropout^35^ layer with a dropout rate of 0.1. Finally, the output layer consisted of an FC layer of *F* units and softmax activation function (here, *F*=3), where each unit encoded either a finger movement or both as not moving. The network was trained using 250 epochs, the categorical cross-entropy as loss function and Adam^36^ as optimizer with a learning rate of 0.01, *β*_1_ as 0.9 and *β*_2_ as 0.999. We trained one model on the OPM signal and one on the EMG signal. Model evaluation was performed with a stratified 5-fold cross-validation by computing the accuracy of the model between the ground truth and its predictions. Notably, we z-scored the data by computing the moments (sample mean and standard deviation computed on the sampled windows dimension) on the train set and then applying them to the test set to avoid possible confounders.

### Feature importance analysis

We investigated how the models generated predictions in the test set by exploiting a recent approach in the field of explainable deep learning^26^, namely the integrated gradients^27^, that allowed us to perform a feature importance analysis. For each sample of the test set, we first linearly interpolated the sample (i.e., a *T* × *C* matrix) with a “baseline” matrix of zeros with the same dimensionality, using 6 levels of transparency linearly sampled from 0 to 1. We passed these interpolated samples to the model and compute the partial derivative (i.e., the gradients) of the loss function with respect to the input. Next, we combined these gradients by computing a numerical approximation of their integral over the interpolated samples using the Riemann sum approximation and normalized them to make sure they were in the same scale. We averaged these values across the temporal dimension of the input and across the samples of the test set to obtain a single value for each fold of the crossvalidation scheme and for each channel.

### Inter-rate reliability analysis

We assessed the consistency of the OPM-MMG and EMG model predictions by computing two metrics of inter-rater reliability. The first one was the percentage agreement, which simply quantifies the percentage of predictions for which both models predicted the same class. The second metric was Cohen’s Kappa, defined as the probability of agreement between the two models normalized by probability of agreement expected by chance.

### Statistical analysis

Statistical analyses were conducted on the comparison between the accuracy values of the two models against the empirical chance level, separately for each model. The empirical chance level was computed using a permutation test approach, by permuting the labels of the train set and repeating the cross-validation for 1000 times to obtain a null distribution. P-value was obtained as the number of values found in the null distribution that exceeded the observed value, while the effect size (Cohen’s *d*) was computed as the difference between the observed value and the mean of the null distribution, divided by its standard deviation. We also compared the percentage agreement values, the Cohen’s Kappa values and the integrated gradients values against the resulting null distribution as above. For the direct comparison between the accuracy of the EMG and OPM models, we ran two tests specifically suited for comparing the performance between two classifiers^37,38^. First, we used the 5*×*2 cross-validation F-test by repeating 5 times a 2-fold cross validation and testing both models on the same data. Thus, we computed the pseudo f-statistic and the p-value using an F distribution with 10 and 5 degrees of freedom^38^. Finally, we also compared the models’ performance using the McNemar test, by computing a 2 by 2 confusion matrices between the models’ predictions. Then, we computed the McNemar statistic and the p-value using a *X^2^* distribution with one degree of freedom^37^.

### Data availability

The data that support the findings of this study are available from the corresponding authors upon request.

## Results

We collected data from a single human participant (male, 28 years old). We measured muscle activation on the forearm (Fig. 1A) to detect flexion movements of the index (DII) or little finger (DV). We simultaneously recorded magnetic and electric signals using 4 biaxial OPM sensors and 4 bipolar surface EMG electrodes, respectively (Fig. 1B). Finger movements were simultaneously recorded using a light sensor. In three different sessions, the participant was instructed to flex a finger and then to return to the rest position (open hand). In the first session, the participant flexed the DII finger, in the second the DV finger and in the third both DII and DV, alternatively.

We computed the temporal envelope of 25 to 100 Hz power of all MMG and EMG signals (see methods), aggregated the data from all the three sessions, and computed the time-course of the signal envelope relative to the onset of DII (Fig. 2A) or DV (Fig. 2B) finger motion for each channel and sensor modality. The visual inspection of movement-locked envelopes suggested that both, EMG and MMG captured muscle activity during finger movement and that EMG had a higher signal-to-noise ratio during finger movement relative to the pre-motion baseline. Furthermore, the pattern of results suggested that channels positioned on the ulnar side of the forearm (EMG-2, EMG-4, OPM-2YZ and OPM-4YZ) were measuring signals more during DV motion, while sensors on the radial side (EMG-1, EMG-3, OPM-1YZ and OPM-3YZ) were measuring signals more during DII movement.

**Figure 2.**
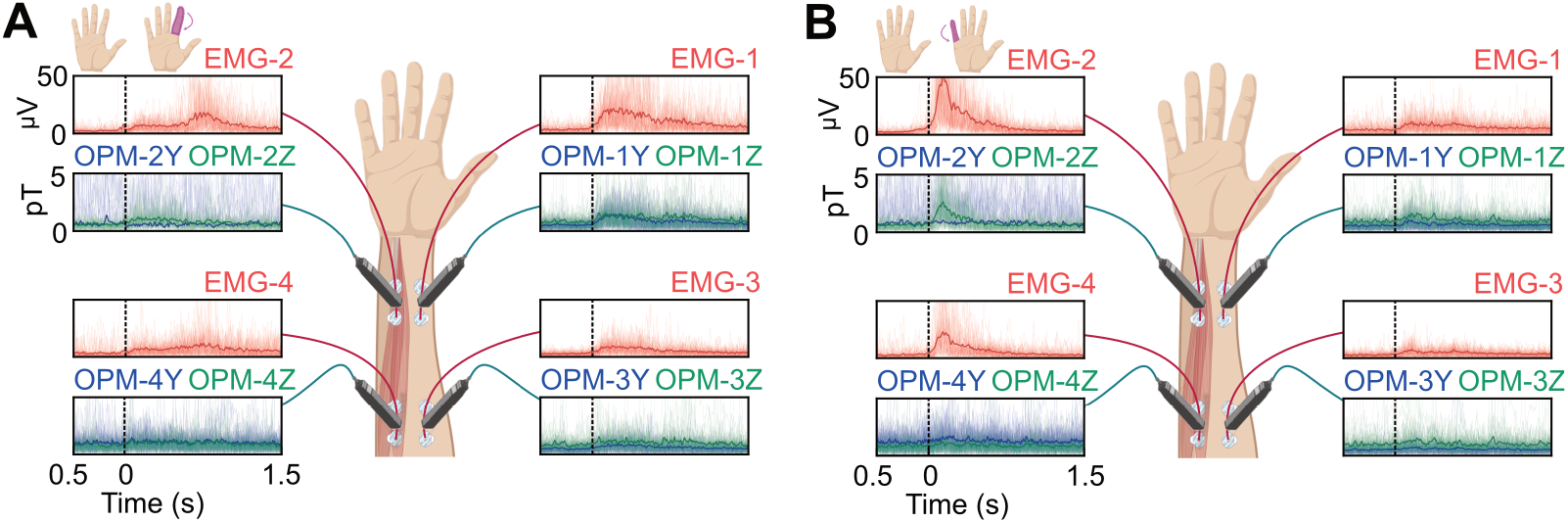
**A.** Line plots showing the time course of the aggregated data of the DII finger movement across the three sessions and divided by channel. **B**. Line plots showing the time course of the aggregated data of the DV finger movement across the three sessions and divided by channel.

### Classification analysis

To quantify these results and to compare the two sensor modalities, we classified finger movements from EMG and MMG signals. Specifically, we performed a multiclass classification analysis on 100 ms temporal windows of either EMG or MMG signals using a Deep Residual Convolutional Neural Network^25^. The 3 classes to predict were whether finger DII was moving, finger DV was moving or both fingers were not moving. We trained two models, one for each sensor modality, using a stratified 5-fold cross-validation scheme with a nested sampling procedure.

We plotted the models’ predictions as a density plot on a 2-dimensional simplex where vertices represented the three classes (Fig. 3A). Visual inspection of these plots showed qualitatively similar distributions across sensor modalities. To assess model convergence, we plotted the loss function (categorical cross-entropy) as a function of the epochs used for training the models for both EMG and MMG models (Fig. 3B). Both models reached their plateau performance around 100 epochs, with the EMG model converging faster than the MMG model.

**Figure 3.**
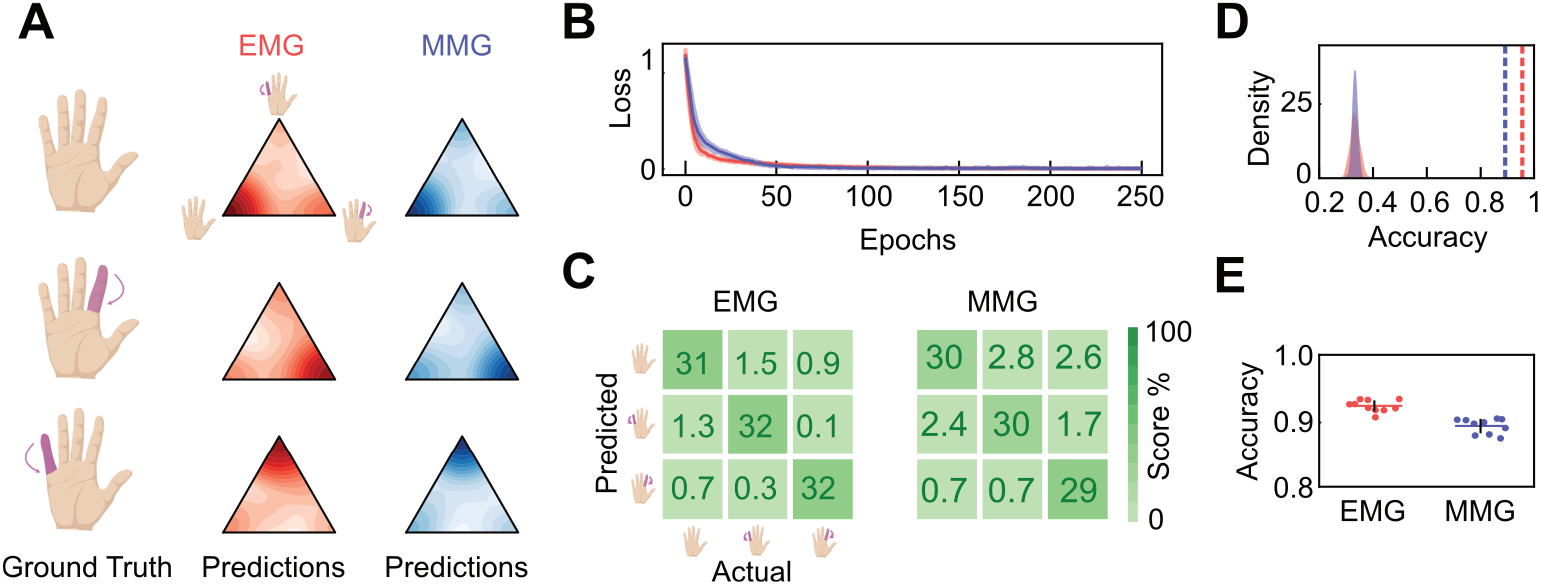
**A.** 2-dimensional simplex density plots of EMG (red) and MMG (blue) models’ predictions divided by classes. **B**. Line plots of the loss function across the epochs for EMG (red) and MMG (blue) models. **C**. Confusion matrices of the EMG and MMG models’ performance. **D**. Results of the permutation test on the accuracy of the EMG (red) and MMG (blue) models. The density plots are the null distributions obtained separately for each model, while the vertical lines are the observed accuracy values, averaged across folds. **E**. Results of the 5*×*2 cross-validation F-test. Scatter dots are individual folds, while error bars represent standard deviation.

To quantitatively assess both models’ performance, we next computed the confusion matrix between the models’ prediction and ground truth (Fig. 3C). Both models showed high-performance in the classification task, with similar patterns of errors and accurate predictions. All three states could be significantly (compared to a null distribution) classified by both models with an overall accuracy of 95.31% (*p* < 0.001, *d* = 34.41, 95% CI [32.89, 35.92]) and 89.06% (*p* < 0.001, *d* = 53.74, 95% CI [51.38, 56.11]) for EMG and MMG, respectively (Fig. 3D). The EMG accuracy was significantly higher than the MMG accuracy (Fig. 3E, *F* (10,5) = 10.66, *p* = 0.009, *d* = 2.97, 95% CI [2.78, 3.16], 5*x*2 cvtest; *X^2^* (1) =9.78, *p* = 0.002, Cohen’s *g* = 0.22, McNemar test).

### Feature importance analysis

After having established the models’ performance, we investigated which channels the models relied mostly on to make predictions. We conducted a feature importance analysis exploiting recent advances in explainable methods in deep learning^26^ such as integrated gradients^27^. We computed integrated gradients for each channel across cross-validation folds and compared them against a null distribution to test their significant contribution to models’ predictions. We found EMG model significantly relied on all channels (Fig. 4A, all *p* < 0.002), even though the channels positioned on the ulnar side of the forearm had a higher effect size (EMG-2 *d* = 79.39, 95% CI [75.91, 82.88], EMG-4 *d* = 24.27, 95% CI [23.21, 25.34]) compared to the others (EMG-1 *d*= 5.03, 95% CI [4.81, 5.26], EMG-3 *d* = 5.28, 95% CI [5.04, 5.52]). For the MMG model, we found that it significantly relied on all channels (all *p* < 0.018) but OPM-1Y, OPM-3Y and OPM-4Y (all *p* > 0.05). We also found that the effect size was generally larger on the ulnar side for the Z-axis (OPM-2Z *d* =18.24, 95% CI [17.43, 19.05], OPM-4Z *d* = 3.66, 95% CI [3.49, 3.84]) compared to the opposite side (OPM-1Z *d* =9.74, 95% CI [9.30, 10.17], OPM-3Z *d* = 2.79, 95% CI [2.65, 2.93]).

**Figure 4.**
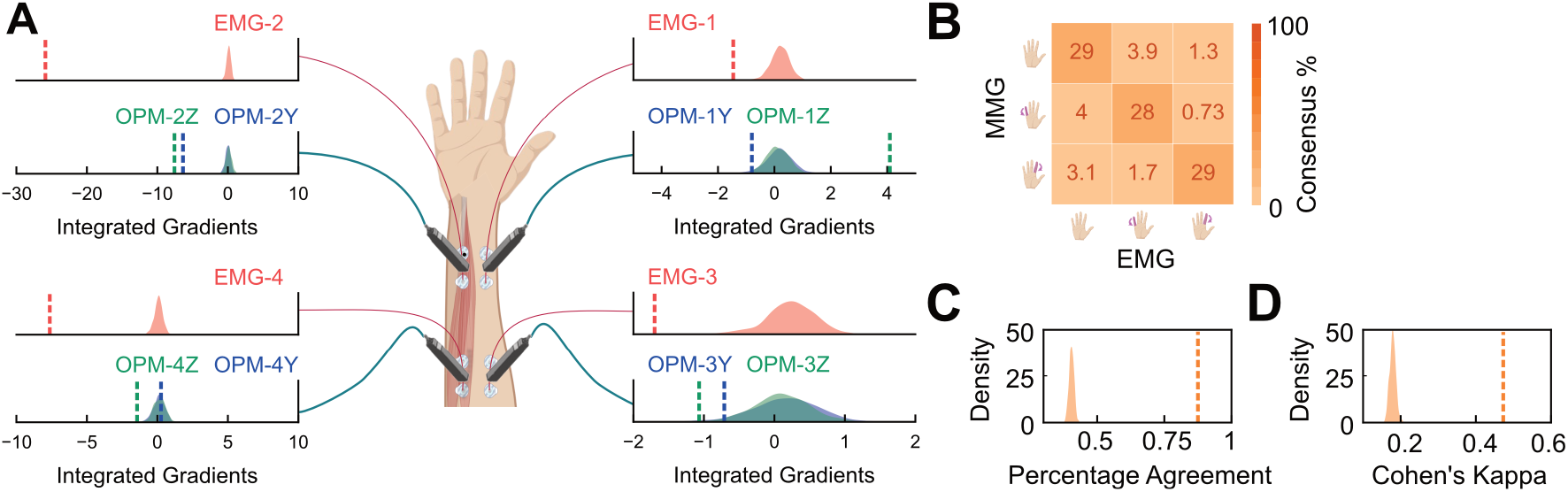
**A.** Results of the permutation test on the integrated gradients values representing the feature importance scores for each channel and sensor modality, i.e. EMG (red) and MMG (blue for y-axis and green for z-axis). The density plots are the null distributions, while the vertical lines are the observed values, averaged across folds. **B**. Consensus matrix between EMG and MMG models’ predictions. **C**. Permutation test showing the percentage agreement values between EMG and MMG models’ predictions. The density plot is the null distribution, while the vertical line represents the observed percentage agreement, averaged across folds. **D**. Permutation test showing the Cohen’s Kappa values between EMG and MMG models’ predictions. The density plot is the null distribution, while the vertical line represents the observed Cohen’s Kappa, averaged across folds.

### Inter-rate reliability analysis

Finally, we investigated the model agreement, by directly comparing their predictions. We computed a consensus matrix, where the row and column entries of the matrix where the MMG and EMG predictions, respectively (Fig. 4B). We found that their predictions were highly aligned. For all three individual states as well as for the average across all states the agreement between MMG and EMG models was significantly higher than expected by chance (Fig. 4C, mean percentage agreement =85.26%, *p* < 0.001, *d* = 48.79, 95% CI [46.65, 50.94]). We also observed that the average Cohen’s kappa of 0.45 between the two models’ predictions was significantly higher than expected by chance (Fig. 4D, *p* < 0.001, *d* = 33.68, 95% CI [32.19, 35.16]).

## Discussion

In this study, we measured muscle activation using OPM-MMG and EMG to detect finger movements. We found that both sensor modalities were able to discriminate DII, DV and nonmovement. Our EMG results add to previous studies showing the capability to discriminate finger movements with EMG ^28–30^ by demonstrating finger movement discrimination using an end-to-end learning framework based on only 4 EMG channels without explicit feature extraction. Our OPM results are, to the best of our knowledge, the first demonstration of finger movement discrimination with OPM-MMG. We show that also for OPM-MMG an end-to-end learning framework can be adopted for efficient movement discrimination.

We found better performance for the EMG model as compared to the OPM-MMG model. This likely reflects a lower signal-to-noise ratio (SNR) of OPM-MMG. On the one hand, this may reflect a genuinely lower sensor SNR. On the other hand, this may be due to geometrical factors. While the surface-EMG was attached to the skin of the forearm, the OPM sensors were positioned above the skin and independently with respect to the forearm. Thus, both the distance between sensors and muscles and their relative motion was larger for OPM-MMG than for EMG.

The feature importance analysis revealed that both sensor modalities relied more on channels positioned on the ulnar side of the forearm to classify finger movements. This can be explained by the fact that the flexor digitorum muscles (DII-DV) are positioned more on the ulnar side of the forearm than the radial side. Notably, the z-axis (perpendicular to the skin) of the OPM sensor, positioned on the ulnar side of the forearm (OPM-2Z and OPM-4Z), was the feature most used by the model. This highlights the relevance of the spatial axis of MMG measurements and suggests that the skin perpendicular axis may be particularly suited to differentiate the signals of finger flexors.

We found that OPM-MMG and EMG models were highly consistent in their prediction of finger movements. This demonstrates the potential of OPM-MMG as an alternative to EMG, since both do not only have similar classification performance, but also consistent prediction patterns. As OPM-MMG allows contactless measurements, it may be particularly suited for clinical applications in which skin contact is undesirable, such as e.g. measurements in autistic patients^31^ or patients with skin-diseases.

Some limitations of this study need to be considered. First, measurements were performed in a single participant. Thus, although the results provide a proof-of-principle and show that the SNR of the OPM-MMG sufficient for use in single subjects, further studies with larger samples are required to validate the results and estimate population variance. Second, we placed sensors only on the ventral forearm because we focused on palmar flexion movements of the fingers. Future studies may add sensors on the dorsal forearm to also exploit extensor muscles’ signal. Lastly, we collected all data in ideal conditions, for example by using a cast to allow only the fingers of our interest to move. Further studies are required to compare sensor modalities in more naturalistic settings, such as free finger movements.

In sum, our findings show that OPM sensors can be employed to reliably discriminate finger movements and incentivize future applications of OPM in magnetomyography.

